# Investigation of potential hinge region for *Mycobacterium tuberculosis* topoisomerase I conformational change during catalysis

**DOI:** 10.64898/2026.02.04.703551

**Authors:** Shomita Ferdous, Yasir Mamun, Thirunavukkarasu Annamalai, Fenfei Leng, Prem Chapagain, Yuk-Ching Tse-Dinh

## Abstract

*Mycobacterium tuberculosis* topoisomerase I (MtbTOP1) is essential for the viability of the causative agent of TB. There are still significant unanswered questions regarding the dynamic conformations during catalysis of relaxation of negatively supercoiled DNA by MtbTOP1. We aim to study the flexible hinge residues that control the dynamics of inter-domain rearrangements involved in the enzyme conformational changes that allow the opening-closing of the topoisomerase gate. We used the online server PACKMAN to predict possible hinges from the MtbTOP1 crystal structure. The predicted region “PRO506 to LEU526” at the border between domains D2 and D4 with a p-value <0.05 was then studied as a potential hinge. The highly conserved ARG516 from this region interacts with the DNA inside the protein toroidal cavity. This arginine maintains inter-domain interaction with GLU207 of D4 and ASP691 of D5 domains. After introducing alanine substitutions, we further studied the mutant topoisomerases in biochemical experiments. The results showed a significant loss in DNA relaxation activity without affecting DNA binding and cleavage after mutating GLU207 and ARG516, consistent with their role as hinge residues in domain rearrangements.

## INTRODUCTION

Type IA topoisomerases are ubiquitous in all domains of life [1,2] and can catalyze topological changes for both DNA and RNA during vital cellular processes [3,4]. This subfamily of topoisomerases includes bacterial topoisomerase I enzymes responsible for relaxing negatively supercoiled DNA to prevent the stabilization of R-loops during transcription elongation [5,6]. Topoisomerase III in prokaryotes and eukaryotes can resolve certain topological barriers generated in replication and recombination [1,2]. In the topological changes catalyzed by topoisomerase IA, a DNA-gate is generated by enzyme cleavage on a single-stranded G (gate) segment coupled to protein conformational change to allow movement of either single-stranded or double-stranded T (transport) segment across the gate to change the DNA topology following religation of the G segment break and release of the DNA.

Based on the available crystal structures of bacterial topoisomerase I [7-11], it has been proposed that the domains D1-D4 from the conserved N-terminal domains (NTD) of topoisomerase IA have to change their relative positions to each other during each catalytic cycle (Figure S1) to adopt open and closed conformations required for entry of the T segment DNA into the interior of the toroidal cavity, followed by religation of the DNA break on the G segment [12]. Previous single molecule studies have shown that the DNA gate formed by *Escherichia coli* topoisomerase IA can open by up to 6.6 nm [13]. These inter-domain relative movements are likely to be achieved through hinge motions. Hinge motions in protein mean rotations of one domain against another around a line separating two planes. According to the MolMovDB database [14], hinge motions account for 45% of all protein collective motions [15]. The overall protein motion is a complex motion combining hinge and shear motions. Since studying protein motions by MD simulations is both financially challenging and time consuming, we need an alternative approach to predict the motions successfully. Deciphering hinge residues and related motion will result in a deeper understanding of protein function and dynamics. The structure-based drug design of protein ligands is mostly based on protein crystal structure, a snapshot of a single conformation. Meanwhile, proteins have multiple conformations resulting from the protein’s motions, and their functions are very closely related to these large conformational changes [16]. Thus, additional knowledge on protein motion will extend our understanding of function-related dynamics of proteins, eventually aiding in design of drugs that can bind to different sites on the topoisomerase target.

The hinge regions are distinct from the rest of the proteins due to the overall packing of amino acid residues and constraints in the hinge regions. Generally, hinge region residues are more flexible and show a lower constraint on amino acid movement than the rest of the rigidly packed domains. The changes associated with hinge motion include changes in protein backbone torsion angle, hydrogen bond, and packing interaction across the hinge [16]. Since hinge regions are more flexible, they can easily change conformation and initiate domain rearrangements needed for the catalysis of topoisomerase IA.

The Online server “PACKMAN” [16] can predict the hinges from either an open or closed form based on protein packing, graph theory, and statistical approaches. Local, flexible regions are detected as potential hinges and validated through permutation test statistics or p-value [16]. We employed PACKMAN in this study to predict possible hinges in the crystal structure of *Mycobacterium tuberculosis* topoisomerase I (MtbTOP1) with DNA bound both as potential G segment and T segment in the structure [11]. MtbTOP1 is essential [17,18] and a validated target for TB drugs [18,19]. A selected hinge region was further investigated by molecular dynamics (MD) simulation and site-directed mutagenesis to confirm its potential role in domain rearrangements required for the relaxation of supercoiled DNA.

## RESULTS AND DISCUSSION

### PACKMAN predicted potential hinge regions at different locations

Crystal structure (pdb_00008czq) of MtbTOP1-704t (truncated version terminating after residue 704 out of 934 total residues) with N-terminal domains D1-D4 plus C-terminal domain D5 was used in PACKMAN to predict potential hinge regions after removing the bound DNA segments from the structure. The online server PACKMAN predicted four different regions of MtbTOP1 as potential hinges, among which three of them have a statistically significant p-value (<0.05). Those regions are “Q680 to T704” from the D5 domain, “P506 to L526” from the D4 and D2 domain interface, and “R250 to D273” from the D2 domains (Figure 1). The first predicted hinge is entirely in the D5 domain, with some polar contacts with domains D2 and D4. Within the next predicted region, P506 to L526, there are two highly conserved polar residues in bacterial topoisomerase I sequences, E519 and R516 as shown by sequence alignment of topoisomerase I sequences from 23 bacterial species (Figure S2). Residue E519 from D4 has been shown to have interdomain interactions with D3 in the available crystal structure [11]. The other polar residue, R516 from D2, interacts with D691 from the D5 domain of the C-terminal domains and E207 from the D4 domain (Figure 1 close-up view). This predicted hinge region resides at the border between D2 and D4 and has parts of an alpha helix, beta-sheet, and disordered loop region (shown in green in Figure 1). The third predicted hinge R250 to D273 from the D2 domain does not show inter-domain polar contacts, but the residue R255 from this region interacts with the negatively charged backbone of the T-segment DNA.

**Figure 1.**
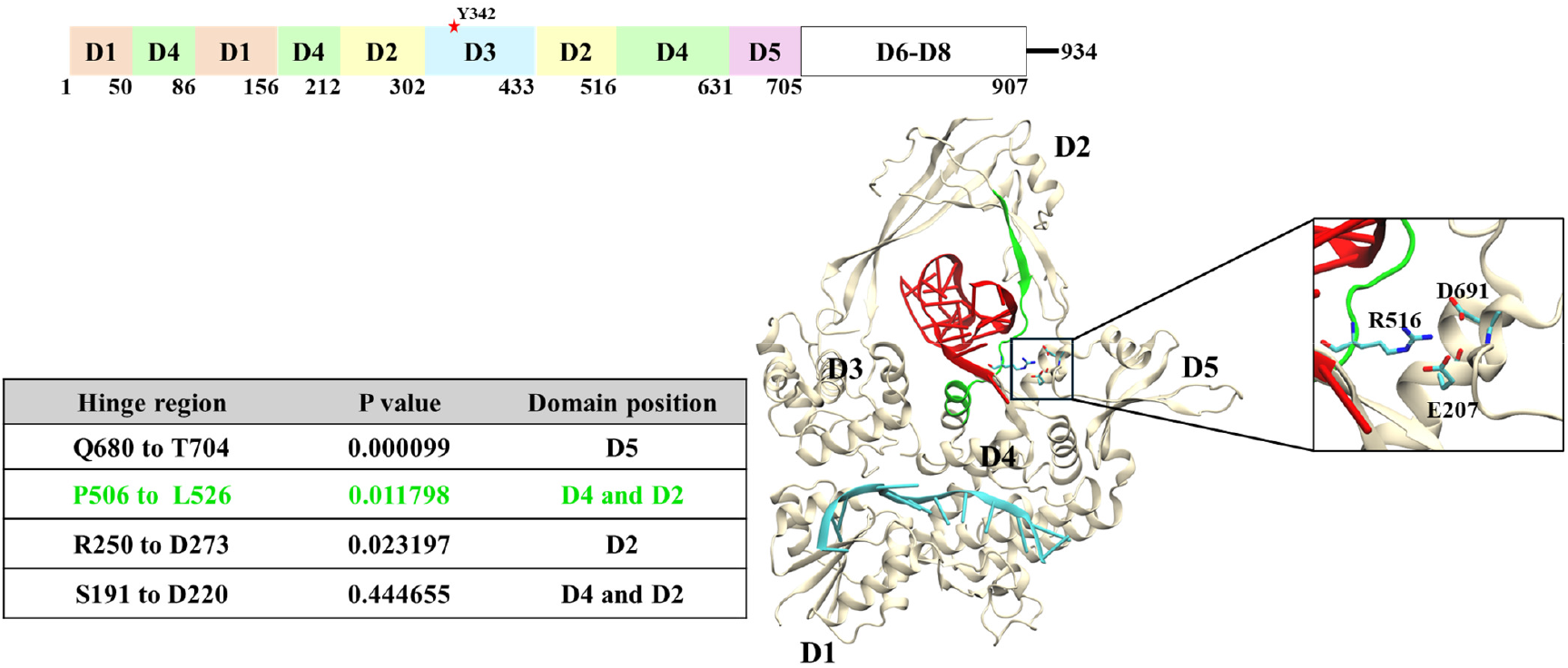
Prediction of hinge regions in MtbTOP1 by PACKMAN. Domain organization of MtbTOP1 and crystal structure (pdb_00008czq) of domains D1-D5 of MtbTOP1 with DNA proposed [11] to correspond to G-segment (colored teal) or T-segment (colored red) are shown. The packman predicted hinge regions of MtbTOP1 are listed with respective p-values. The predicted hinge region from P506 to L526 is colored in green. The interactions between residues R516, E207 and D691 are shown in the close-up view.

### Molecular dynamics (MD) simulation of the interaction between T-segment DNA and the predicted hinge regions

Following the PACKMAN prediction, we further investigated the predicted hinge region from P506 to L526 because of its potential for facilitating the movement of domains D2, D3 away from domain D1 to open the topoisomerase gate We ran a 500 ns MD simulation on structure 8CZQ (pdb_00008czq) [11] which has a G-segment DNA bound at the active site and also a duplex DNA postulated to correspond to the T-segment DNA present inside the toroidal cavity of MtbTOP1 to provide the maximum structural information on the potential hinge region (Figure 2A). When we measured the H-bond interactions between the target residues themselves and with the T-segment DNA, we observed that residue R516 showed very strong polar contacts or H-bonding with a Guanine (Table 1, Figure 2A), maintaining a close proximity during the length of the simulations (Table 1, Figure 2B). This suggests a potential role for R516 in the entry and retention of T-segment DNA during catalysis. Previously, residues in close proximity to the hinge region in both archaeal *Caldiarchaeum subterraneum* and human topoisomerase IB [20] have been proposed to influence DNA entry and interaction. We hypothesize that residue R516 could play a similar role in the hinge region of MtbTOP1.

**Table 1.**
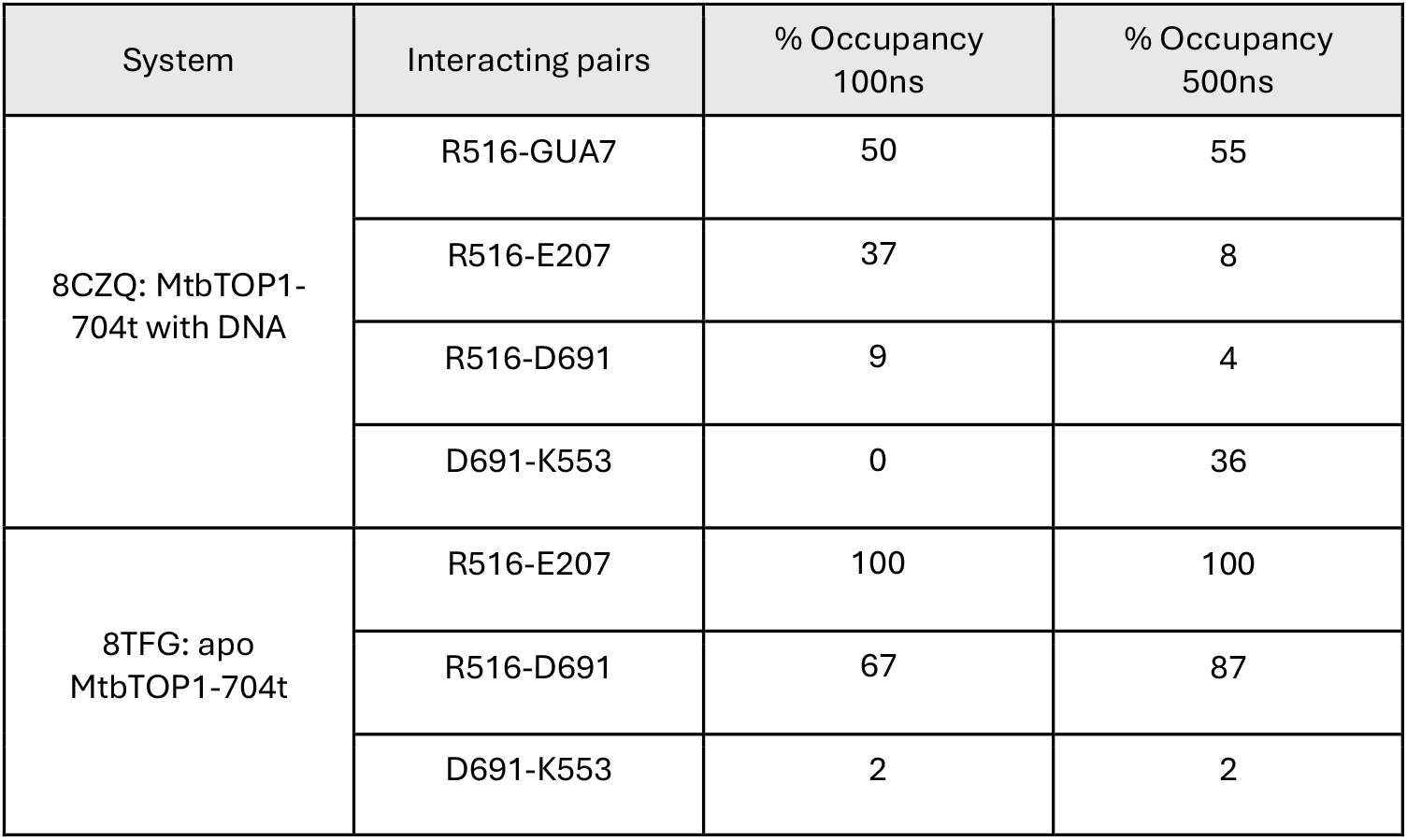
Percent H-bond occupancy between residues or with T-segment duplex DNA in the systems after molecular dynamics simulation.

| System | Interacting pairs | % Occupancy<br>100ns | % Occupancy<br>500ns |
| --- | --- | --- | --- |
| 8CZQ: MtbTOP1-704t with DNA | R516-GUA7 | 50 | 55 |
|  | R516-E207 | 37 | 8 |
|  | R516-D691 | 9 | 4 |
|  | D691-K553 | 0 | 36 |
| 8TFG: apo MtbTOP1-704t | R516-E207 | 100 | 100 |
|  | R516-D691 | 67 | 87 |
|  | D691-K553 | 2 | 2 |

**Figure 2.**
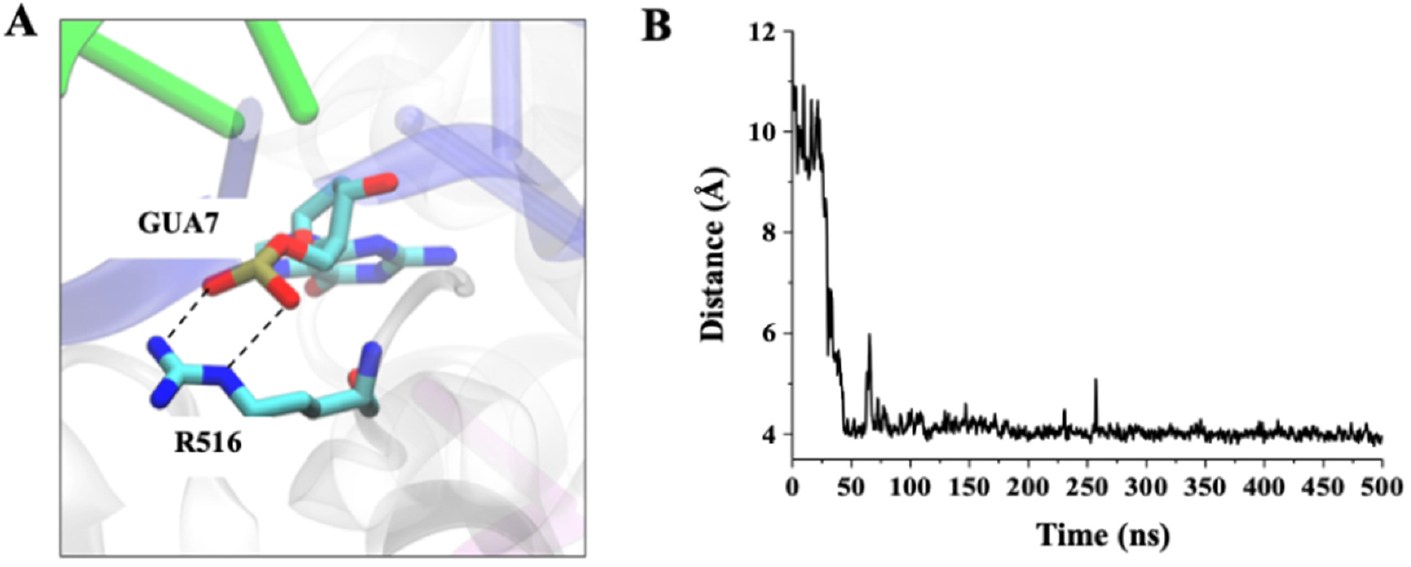
Interaction between a proposed hinge residue and T-segment DNA. A) Close-up figure showing the interaction between residue R516 and the 7^th^ base (GUA7) of one strand of the DNA bound within the toroid hole of MtbTOP1. B) Distance plot highlighting the proximity of R516 and GUA7 during the length of the simulation.

### Inter-domain interactions shared by residue R516 from the predicted hinge region

To compare the inter-domain interactions of residues at the predicted hinge region in the absence of the T-segment DNA, we ran another 500 ns MD simulation for the crystal structure 8TFG (pdb_00008tfg) corresponding to the apo-enzyme. For the 8CZQ system, although R516 shows interdomain polar contact with E207 and D691 in the beginning (37% for E207 and 9% for D691 after the first 100 ns, Table 1), these interactions get weaker as the simulation run continues (8% for E207 and 4% for D691 after 500 ns, Table 1, Figure 3B). However, we observe a stronger interaction between residues K553 and D691 at 500 ns. This contrasts with the 8TFG system, where the interaction between R516 and E207, as well as D691, is strong at both 100ns and 500ns, but the interaction between D691 and K553 is weak overall. (Table 1, Figure 3B). Distance analysis between these residues follows the same pattern as the H-bond interactions (Figure 3C). In the apo 8TFG model, the distances between R516 and E207 and between R516 and D691 remain close, with some fluctuations, while the residues K553 and D691 remain distant. In the 8CZQ system, however, the distance between R516 and E207 and D691 is not as close as in the 8TFG system, but the distance between K553 and D691 becomes very close after a while and remains that way for a long period of time. (Figure 3C).

**Figure 3:**
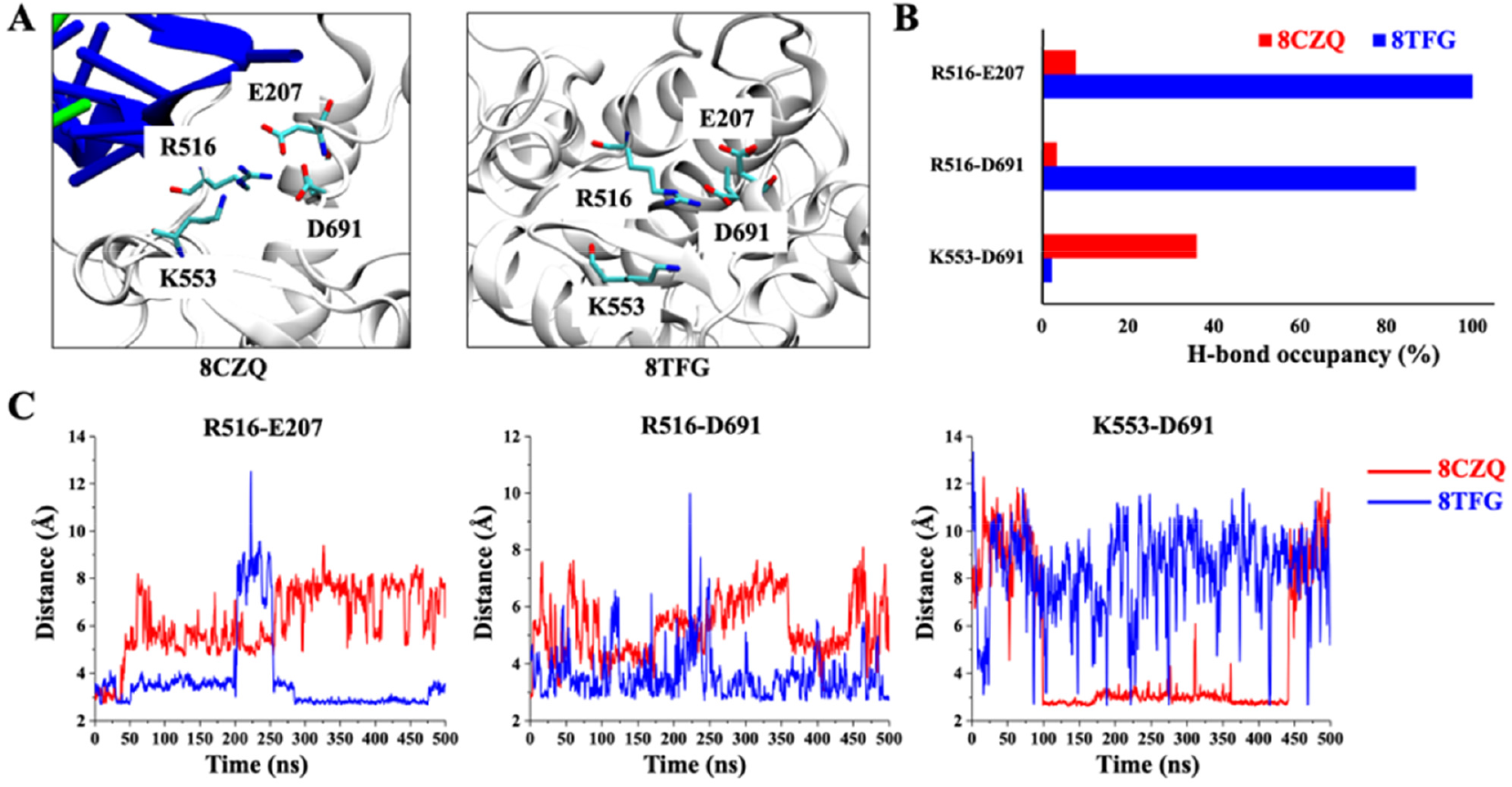
Position and interactions of residues observed from MD simulations. A) Relative positions of hinge region residues R516, E207, K553 and D691. The residues are shown in the stick representation, while protein and DNA (where present) are shown in the cartoon representation. B) Bar plot diagram showing the H-bond occupancy percentage between residues. The values indicate the H-bond occupancy (%) determined using VMD. The occupancy (%) was calculated with a 3.5 Å distance cut-off and a 30° angle cut-off. C) Plot showing the distance between the interacting residues in the 8CZQ (red) and 8TFG (blue) models for the total length of the runs. Distance was measured between the sidechains of the residues.

### Effect of predicted hinge region mutations on MtbTOP1 relaxation activity

Based on the results from the MD simulations, we selected residues R516 in D2 domain and its two potential interacting partners, D691 and E207 for further investigation. The interdomain interactions involving these three residues could potentially affect the relative arrangements of domains D2, D4 and D5 for conformational change during catalysis. Alanine substitution at these three residues were introduced by site-directed mutagenesis. T518 near R516 was also mutated to alanine for comparison. Assay of the relaxation activity using serial dilutions of the wild-type and mutant enzymes showed ∼16-fold and 8-fold lower activity for the R516A and E207A mutant enzyme respectively (Figure 4A). The D619A mutant showed only 2-fold lower activity than wild-type, while the T518A mutant showed no decrease in relaxation activity.

**Figure 4:**
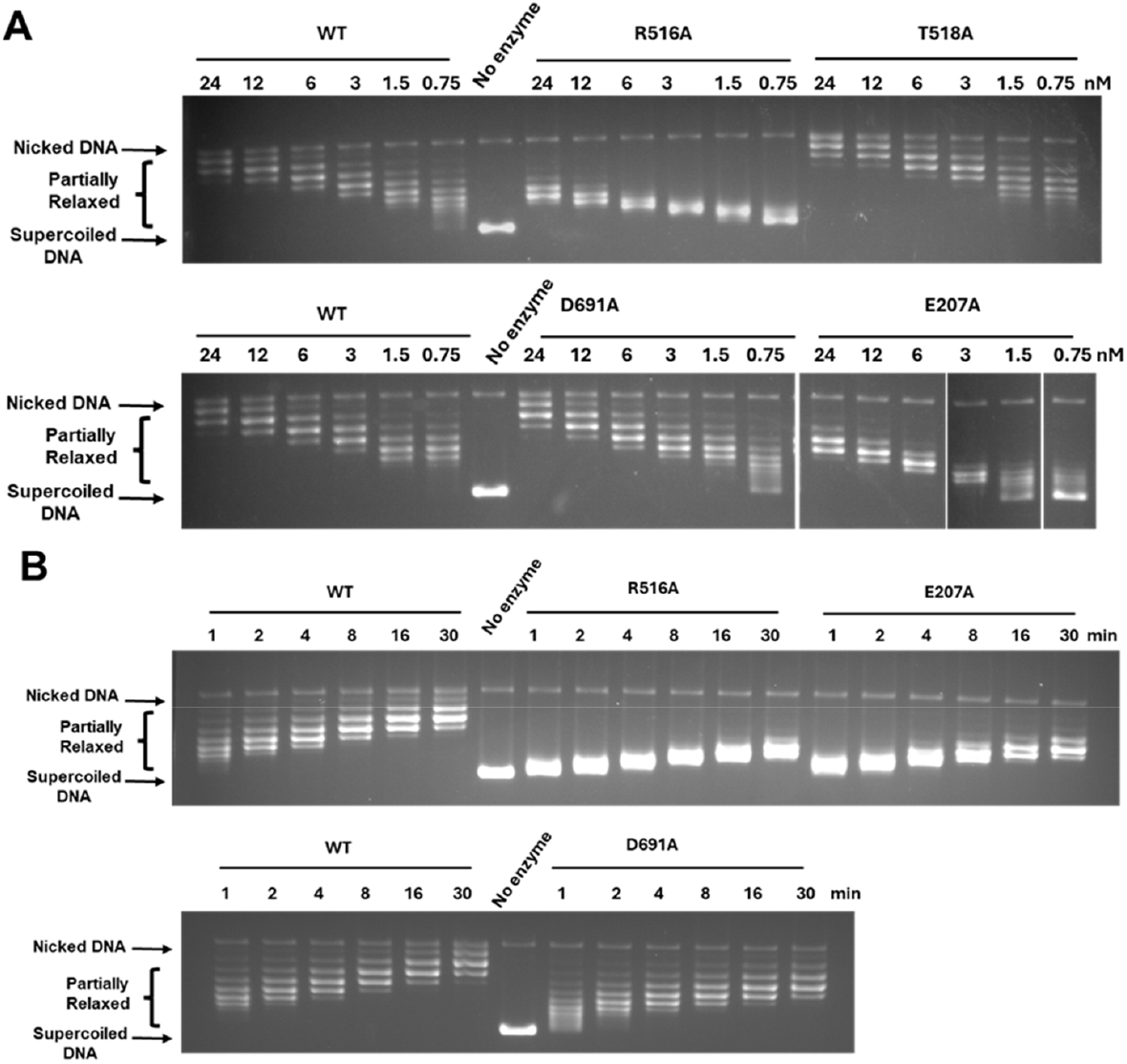
Assay of effect of alanine substitutions at hinge region on relaxation activity of MtbTOP1. A) Relaxation of negatively supercoiled DNA by wild-type and mutant MtbTOP1. Supercoiled pBADthio plasmid was incubated with serial dilutions of enzyme at 37° C for 30 min. B) Time course of relaxation of negatively supercoiled DNA by 10 nM of wild-type and mutant MtbTOP1.

Time course assays were also performed to compare relaxation efficiencies of the mutants at 37°C (Figure 4B). Here, we clearly observed that with the wild-type enzyme, relaxed topoisomers can be observed within 1 minute and the removal of negative supercoils is nearly complete after the first 10 minutes. Whereas, both R516A and E207A showed a delay in producing the relaxed topoisomers, and the relaxation reaction is not yet complete at 30 minuets (Figure 4B). The results from D691A are in agreement with the results from the serial dilution, and showing nearly equal efficiency as the wild-type enzyme (Figure 4B).

### DNA cleavage activity is not affected by the predicted hinge region mutations

According to our hypothesis, the cleavage activity should not be affected significantly after mutating R516, D691, or E207 to alanine since all these residues are far from the cleavage site and are proposed to affect steps in the catalytic cycle that follow the cleavage of G-strand DNA. We used 5’-^32^P-labeled 32-mer oligonucleotide STS32 with a preferred MtbTOP1 cleavage site [21] as the substrate. The results from measuring DNA cleavage extent revealed very similar degrees of cleavage by wild-type and single mutant enzymes at varying enzyme concentrations of 400, 200, 100, and 50 nM (Figure 5A,B).

**Figure 5.**
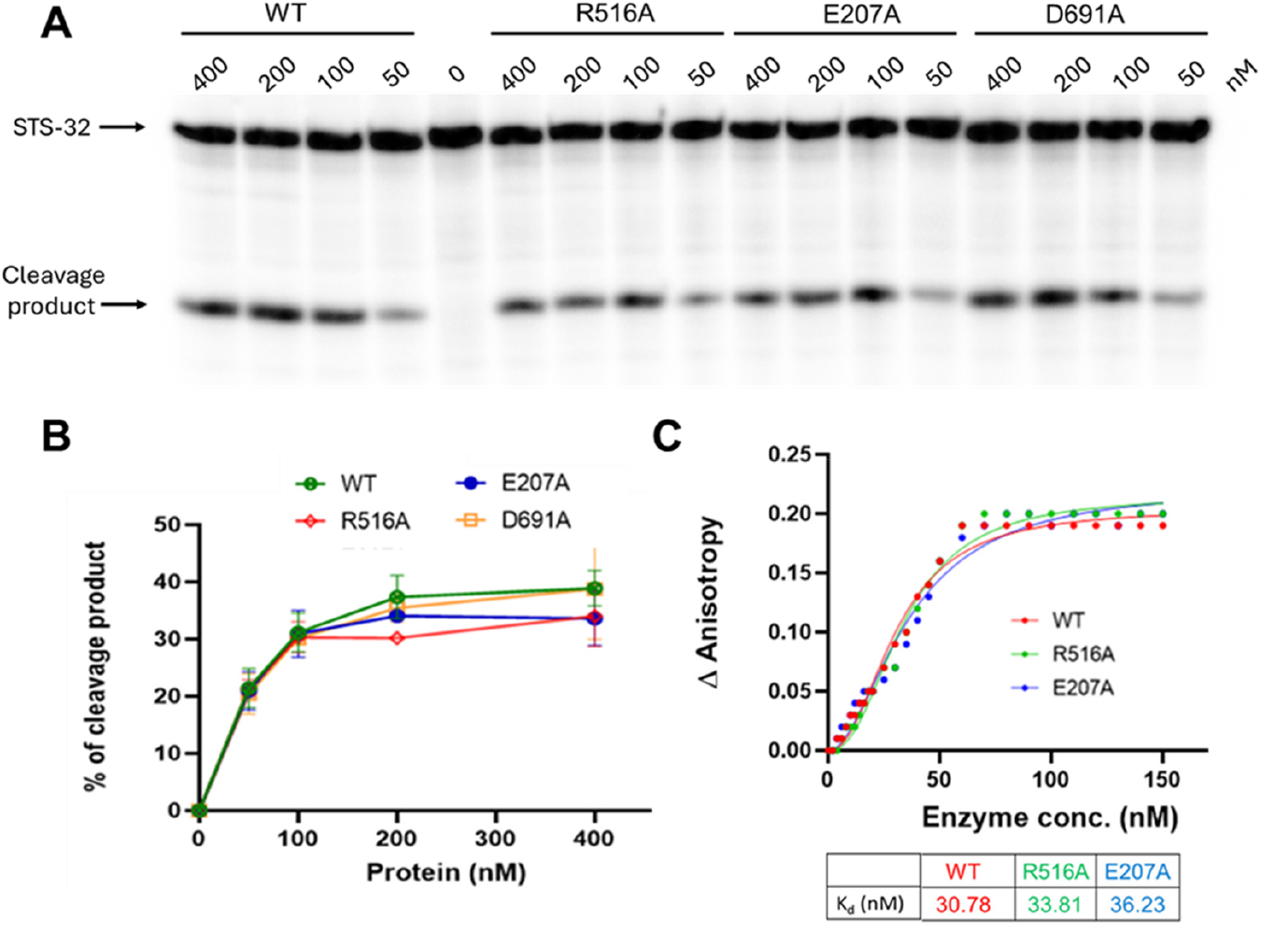
ssDNA cleavage and binding by wild-type and mutant MtbTOP1. A) DNA Cleavage activity shown on 15% denaturing polyacrylamide gel. 100 nM of STS32 oligonucleotide labeled with ^32^P at the 5’-end was incubated for 30 min at 37° C with indicated amounts (nM) of wild-type and mutant MtbTOP1. B) % cleavage quantification for the wild type and mutant MtbTOP1. C) ssDNA binding affinity of wild-type and mutant MtbTOP1 measured by the anisotropy assay.

### ssDNA binds the predicted hinge residue mutants with equal affinity as wild-type MtbTOP1

To measure the ssDNA binding affinity, we incubated a 5’-Fam labeled STS32 oligo with increasing concentrations of wild-type, R516A, and E207A mutant enzymes at room temperature and measured the change in fluorescence signal and anisotropy. The data obtained were further processed in GraphPad Prism to obtain the dissociation constant Kd from the hill fit equation [22]. The Kd values as measurement of the binding affinity of the mutants R516A, E207A, and the wild-type MtbTOP1 are 33.81, 36.23, and 30.78 nM of Kd, respectively, indicating approximately equal binding affinity for ssDNA (Figure 5).

## DISCUSSIONS

The PACKMAN server used in our study can reliably predict the potential flexible hinge from only one conformation, making it particularly suitable for intrinsically flexible bacterial topoisomerase I that has been crystallized only with the N-terminal catalytic domains bound to each other in the “closed” conformation. To include potential effect of DNA binding on the protein conformational changes that may occur during catalysis, we used as input an *M. tuberculosis* topoisomerase I crystal structure which has both G-segment and T-segment DNA bound (PDB 8CZQ) (Figure 1). The enzyme interacts primarily with only one strand of the duplex DNA bound within the toroid hole, so the same interactions can occur with a single-stranded DNA as T-segment as expected for the relaxation of negatively supercoiled DNA (Figure S1). We have proposed that the 8CZQ structure represents a snapshot of conformation right after re-sealing the cleaved G-segment, followed by gate closing to entrap the T-segment [11]. It is notable that all the predicted hinge outputs from using 8CZQ as an input structure are at the right rim or the side of the D2-D4-D5 interface. There is still the possibility that additional hinges may be located at different sites or close to the interface of D3 and D4 at the left rim for conformational change that takes place linked to the initial gate opening in the catalytic cycle.

From the three predicted hinges with statistically significant *p-*values, we decided to investigate “P506 to L526” first because of its location at the D2-D4 interface and having a loop region, which imparts flexibility or movement to this region (Figure 1). This region also consists of highly conserved polar residues like R516, T518, and E519, which participate in electrostatic and hydrogen bond interactions with other residues and maintain inter-domain connections. E519 in D4 can form a strong electrostatic interaction with the positively charged terminus of an alpha helix in adjacent D3, and this interaction is diminished when D4 and D3 move away from each other upon DNA binding [11]. The residue R516 is promising as a hinge residue since it has been found to participate in multiple interactions among D2, D4, and D5 domains and can potentially function as a signaling residue between NTD and CTD, as well as a switch for gate dynamics. Moreover, the interaction between R516 and T-segment DNA bound in the central cavity is one of the strongest interactions between the MtbTOP1 amino acids and T-segment DNA obtained from analyzing the frames of a 500 ns MD simulation (Figure 2). R516 can potentially be a residue that can initiate a conformational change upon DNA binding, playing a similar role as the conserved tyrosine adjacent to the hinge of topoisomerase IB [20]. Our hypothesis is supported by the in vitro findings on the alanine substitution mutation of R516. The relaxation activity of the R516A mutant topoisomerase is reduced by ∼16-fold (Figure 4). The close to wild-type ssDNA binding and cleavage activity for the R516A mutant further supports the hypothesis that R516 has roles in gate dynamics and T-segment DNA interaction that follow G-segment binding and cleavage (Figure S1).

The results from the alanine substitution mutant of the R516 interacting partner E207 also support our hypothesis. E207A mutant has a reduction in relaxation activity as the R516A mutant (Figure 4). Looking at the relaxed topoisomer formation during the relaxation assay, we can see that the number of topoisomers produced by the E207A and R516A mutants is fewer than that of the wild-type enzyme. This set of data thus indicates that, due to the R516 and E207 mutations, the MtbTOP1 has lower enzyme processivity with fewer negative supercoils removed before the enzyme dissociates from the plasmid DNA substrate. The relaxation time-course assay at 37° C (Figure 4B), R516A and E207A showed that both mutants have reduced relaxation efficiency compared to WT. The cleavage product formation by E207A is also similar to wild-type MtbTOP1 (Figure 5). Since E207A is not taking part in T-segment DNA interaction, the decrease in relaxation activity for the E207A mutant is probably due to effect on the enzyme conformational change during the catalytic cycle. We did not see any large effect from the D691A substitution on either the relaxation or cleavage activity of MtbTOP1 (Figures 4 and 5). This could be because a negatively charged glutamate E692 is proximal to D691 to potentially compensate for the negative charge loss for interaction with R516 (Figure S3).

We investigated if the identified hinge region or the residues directly interacting with the hinge region of the MtbTOP1 can be a possible drug target with multiple algorithms that can identify drug binding pockets in proteins. The high-resolution crystal structure of MtbTOP1 (8TFG) was uploaded to the online servers DoGsitescorer [23]and CavityPlus 2022 [24]. While the methods of operation for these servers differ, the results generated are comparable, as they all provide a visual depiction of the volume, druggability, and neighboring residues of the predicted drug-binding pocket. Both servers identified possible drug binding pockets in the structure. From the CavityPlus 2022 prediction, a drug binding pocket sharing residues from multiple domains, including C-terminal domain residues, showed a higher binding score than the active site for G-segment cleavage (Figure 6A, Table S3). This predicted binding site contains the residue E207 from D4 and D691 from D5 domain which directly interact with the hinge region studied here. The other server, DoGsitescorer, also shows a binding pocket with a high drug score, consisting of residues from D4 and D5 domains (Figure 6B, Table S4). The best drug binding pocket, according to all the servers, was the G-segment DNA binding cavity in the N-terminal domain D4, possibly due to the large volume of the cavity (Tables S3, S4).

**Figure 6.**
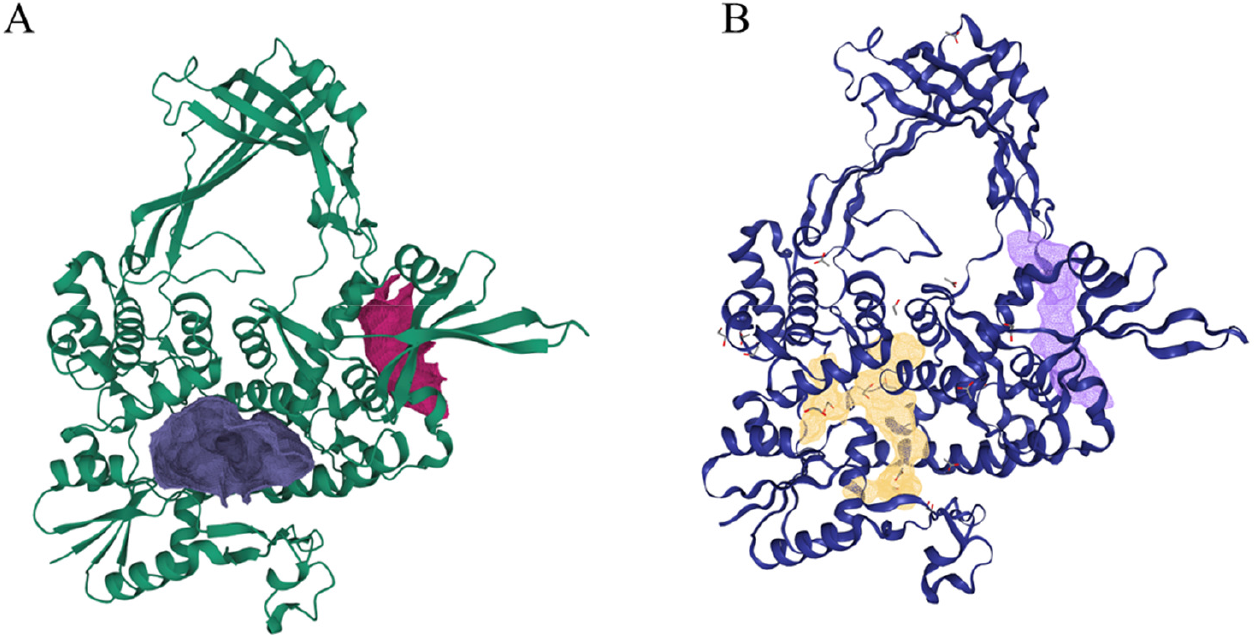
Identification of possible drug binding pockets in MtbTOP1. Depiction of possible drug binding pocket at the DNA binding cavity and hinge region identified by CavityPlus 2022 (A), and DoGsitescorer (B). The enzyme is shown in the carton representation with the possible pockets are shown in the surface representation. While other pockets are also identified (Table S3, S4), only two are shown here for clarity.

## CONCLUSIONS

In summary, we propose a model highlighting the crucial hinge region residues and their roles in gate dynamics. Results from site-directed mutants support a role for R516 in inter-domain networking and T-segment DNA interaction for the relative movement of domain D3 away from D4 and D1 during catalysis. The E207 residue is an interacting partner of R516 that may also participate in gate dynamics. In future, we aim to study the other predicted hinge regions with significant p-value and possibility of drug binding pockets to target putative hinge regions for screening of inhibitors.

## METHODS

### In-silico hinge prediction

For the initial prediction of the flexible hinge region, we employed the online server PACKMAN developed at Iowa State University [16]. The crystal structure 8CZQ, a structure of MtbTOP1 bound to both G and T-segment DNA [11] was used as input for maximum structural information. Before applying 8CZQ as the input for PACKMAN, we rebuilt the missing residues using Modeller [2**5**]. We narrowed down the predicted hinge regions based on the statistically significant p-values (p<0.05).

### Site-directed mutagenesis

Wild type and mutant MtbTOP1 gene were inserted in a previously described clone [26]derived from pET-His6-Mocr TEV LIC cloning vector (2O-T, Addgene plasmid #29710) having a His6 Mocr (monomeric mutant of OCR from bacteriophage T7) tag. The desired MtbTOP1 mutations were introduced through site-directed mutagenesis by PCR amplification with the Q5 site-directed mutagenesis kit from New England BioLab following the protocol provided. The primers for each mutant were designed on the NEBasechanger website, and the sequences are listed in Table S1. The mutant plasmids were cloned into NEB 5-alpha cells for isolation. Complete sequence of mutant plasmids was confirmed by Plasmidsaurus sequencing service through long-read sequencing technology.

### Protein expression and purification

The mutant and wild-type plasmids were transformed into *E. coli* C41(DE3) strain (from Lucigen). The transformants were grown overnight at 37°C in LB (Miller) broth with 100 μg/mL carbenicillin and 1% glucose, followed by a 1 to 100 ratio dilution in fresh LB broth with carbenicillin the next morning. The cells were grown at 30°C till OD600 reached 0.4 and kept on ice for 15 min. Next, 1 mM final concentration of IPTG (isopropyl-D-thiogalactopyranoside) was added to the cells for inducing the overexpression of MtbTOP1 by T7 RNA polymerase under the control of lacUV5 promoter. After overnight induction at 20°C, the cells were pelleted and lysed by adding lysis buffer containing 20 mM sodium phosphate pH 7.4, 0.5 M NaCl, 20 mM imidazole, and 1 mg/mL lysozyme and kept on ice for an hour. Four cycles of freeze thaws were then carried out to ensure complete cell lysis. The soluble lysate with the His6-Mocr-tagged protein was collected after centrifugation at 35,000 rpm for 1 h and then mixed with Ni Sepharose Fast Flow resin (GE Healthcare) for 1 h at 4°C. The affinity resin packed into a column was washed extensively with a buffer of 20 mM sodium phosphate, pH 7.4, 0.5 M NaCl, and 20 mM imidazole before elution of the His6-Mocr-tagged protein with a buffer containing 20 mM sodium phosphate, pH 7.4, 0.5 M NaCl and 500 mM imidazole. The eluted His6-Mocr-tagged protein was incubated with tobacco etch virus (TEV) protease (1 mg of TEV protease for every 40 mg of fusion tagged protein) at 20°C for 6 h, followed by 4°C overnight in buffer of 50 mM Tris, pH 8, 0.5 mM EDTA and 1 mM dithiothreitol (DTT). The TEV-digested sample was then passed through the Ni Sepharose again to remove the fusion tags. The MtbTOP1 protein from the flow-through was further purified with S300 size exclusion chromatography (20 mM Tris–HCl, pH 8, 300 mM KCl). The pure fractions were then concentrated and dialyzed into the storage buffer (100 mM potassium phosphate pH 7.4, 1 mM EDTA, and 50% glycerol) for biochemical assays.

### Relaxation Activity Assay

The reaction volume of 20 μL contained 10 mM Tris–HCl, pH 8.0, 50 mM NaCl, 0.1 mg/mL gelatin, 2 mM MgCl_2_, and 0.3 µg of negatively supercoiled pBAD/thio plasmid DNA (5.5 nM) purified by cesium chloride gradient centrifugation. Serial dilutions of wild-type and mutant MtbTOP1 enzymes were added to the reaction at concentrations of 24, 12, 6, 3, 1.5, 0.75 nM followed by incubation at 37°C for 30 min or the indicated length of time. The reactions were stopped by adding 5 μL of 4X stop solution (50 mM EDTA, 50% glycerol, and 0.5% v/v bromophenol blue). The relaxation reaction products were analyzed by electrophoresis in a 1% agarose gel with TAE (40 mM Tris-acetate, pH 8.0, 2 mM EDTA) buffer at 25 V (1 V/cm) for 18 h. The gel was stained for 1 h in 1 µg/mL ethidium bromide solution, destained with deionized water for 15 min, and photographed using UV light with the Alpha Imager Mini.

### Cleavage Assay

A 32-mer long oligonucleotide STS32 (5’-CAGTGAGCGAGCTTCCGCTTGACATCCCAATA-3’) [21] was labeled with ^32^P at the 5’-end with T4 polynucleotide kinase. STS32 has been shown to yield 18-mer long product of 5’-^32^P-CAGTGAGCGAGCTTCCGC-3’ when cleaved by MtbTOP1 [21,26]. Wild-type and mutant MtbTOP1 enzymes were serially diluted in a buffer containing 100 mM potassium phosphate, 0.2 mM EDTA pH 8.0, 50% glycerol, 1.0 mM DTT, pH 7.4 to 400, 2000, 1000, and 500 nM concentrations. 5 μL reaction mix was prepared with 10 mM Tris-HCl, 1mM EDTA, pH 8 and 100 nM ^32^P-labeled STS32 oligonucleotide, and 0.5 μL of serially diluted enzymes for final enzyme concentrations of 400, 200, 100 and 50 nM, respectively. The reactions were incubated at 37°C for 30 min, followed by the addition of 5 μL of 2X stop mix containing 79% formamide, 0.2 M NaOH, and 0.04% bromophenol blue. The reactions were heated at 95°C for 5 min before loading into a 15% sequencing gel and electrophoresed for 3 h at 20 V/cm in TBE running buffer (0.89 M Tris-HCl, 0.89 M Boric acid, and 0.02 M EDTA, pH 8.0). The gel was then exposed overnight to a phosphor screen and analyzed with the Pharos FX Plus Phosphor-Imager (Bio-Rad). The band intensities were quantified with the QuantityOne software.

### Oligo binding anisotropy assay

The ssDNA oligo binding assay was performed on wild-type and mutant MtbTOP1 following the previously published protocol [22,27,28]. The oligonucleotides STS32 with 6-carboxyfluorescein (6-Fam) at the 3’ end was used as substrate in anisotropy measurements in 50 mM Tris–HCl, pH 7.5, 100 mM NaCl, 0.1 mM EDTA at room temperature. A set of control experiments were performed each time by titration with buffer in the same volumes as the proteins added. The anisotropy measurements were recorded using the Varian Cary Eclipse fluorescence spectrophotometer with excitation wavelength set at 495 nm and emission wavelength set at 520 nm using excitation and emission slits of 5 and 10 nm. The obtained data were further processed in GraphPad Prism to calculate the dissociation constant Kd from the hill fit equation [22,28].

### Molecular Dynamics (MD) simulations

The crystal structures were obtained from PDB. Missing residues were rebuilt using Modeller. The structures were prepared for MD simulation using the solution builder tool in CHARMM-GUI [29-31] by solvating in a cubic box with TIP3P water model and 0.15M concentration of NaCl. The systems 8CZQ and 8TFG were simulated with the CHARMM36m force field. The final size of the water box and the number of atoms in each system are given in Table S2. Simulations were performed with the GPU versions of NAMD 3.0 [32]. Ensembled systems were minimized for 10,000 steps and equilibrated for 250 ps at 303.15 K and 1 atm pressure. Langevin temperature coupling with a damping coefficient of 1/ps was used to keep the temperature constant, and the Nose−Hoover Langevin piston [33] with a 50 fs period and 25 fs decay was used to maintain constant pressure. The Particle Mesh Ewald method (PME) [34] was used for long-range electrostatic interactions with periodic boundary conditions and a non-bonded cut-off set at 12 Å, with smoothing switched on at 10 Å. The covalent bonds involving hydrogen atoms were constrained by ShakeH [35]. The trajectories were analyzed with VMD. VMD and ChimeraX [36] were used to visualize and render the images.

## AUTHOR INFORMATION

### Present Address

Shomita Ferdous, Department of Radiation Oncology, Miller School of Medicine, University of Miami, FL, USA..

### Author Contributions

S.F., Y.T. designed research; S.F., Y.M., and T.A. performed research; S.F., Y.M., T.A., F.L., P.C., and Y.T. analyzed data; S.F., Y.M. and Y.T. wrote the paper.

### Notes

The authors declare no competing financial interest.

## ACKNOWLEDGMENT

This work was supported by National Institute of General Medical Sciences of the National Institutes of Health under Award Number R35GM139817 (to Y.T.).

---

**Table S1.**
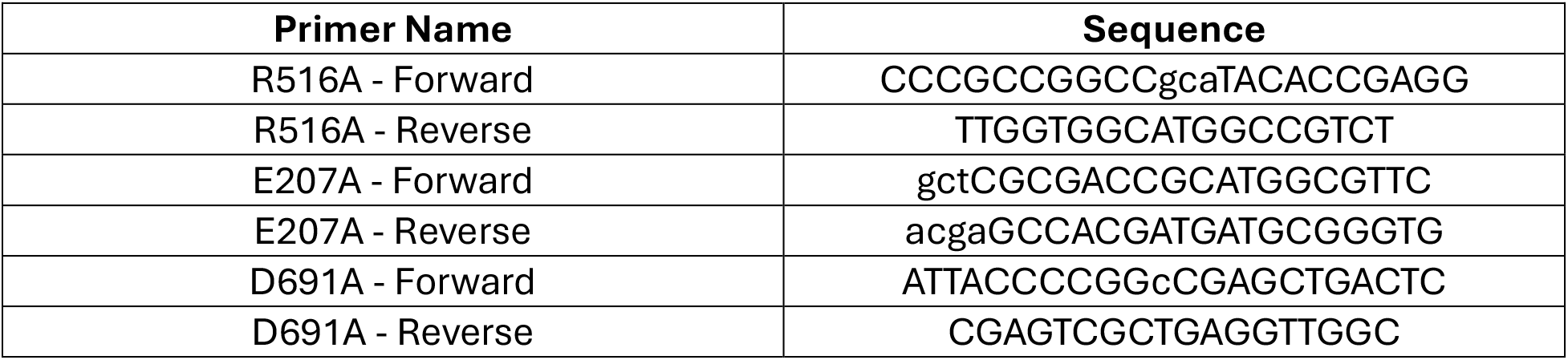
Primers for genertaing MtbTOP1 hinge region mutants.

**Table S2.** Molecular systems simulated in this work. Systems were created using CHARMM GUI and analyzed with VMD.

| MtbTOP1 systems | Ion concentration | Box size ($\text{\AA}^3$) | Atom | Simulation time |
| --- | --- | --- | --- | --- |
| 8CZQ | 0.15 M NaCl | $121 \times 121 \times 121$ | 166,354 | 500 ns |
| 8TFG | 0.15 M NaCl | $122 \times 122 \times 122$ | 179,578 | 500 ns |

**Table S3:** Results from the drug binding pocket prediction webserver CavityPlus 2022.

| Pocket | Pred Max pKd | Pred Ave pkd | DrugScore | Druggability |
| --- | --- | --- | --- | --- |
| 1 | 7.78 | 6.76 | -325 | Undruggable |
| 2 | 10.72 | 6.95 | -340 | Undruggable |
| 3 | 11.18 | 6.45 | -179 | Less druggable |
| 4 | 10.19 | 6.11 | -222 | Undruggable |
| 5 | 9.83 | 5.99 | -638 | Undruggable |
| 6 | 9.43 | 5.85 | -466 | Undruggable |
| 7 | 8.59 | 5.56 | -837 | Undruggable |
| 8 | 7.66 | 5.24 | -837 | Undruggable |
| 9 | 7.44 | 5.17 | -702 | Undruggable |
| 10 | 7.3 | 5.12 | -851 | Undruggable |
| 11 | 7.27 | 5.11 | -699 | Undruggable |
| 12 | 7.23 | 5.1 | -790 | Undruggable |
| 13 | 7.02 | 5.02 | -1094 | Undruggable |
| 14 | 6.91 | 4.99 | -1124 | Undruggable |
| 15 | 6.91 | 4.99 | -886 | Undruggable |
| 16 | 6.75 | 4.93 | -753 | Undruggable |
| 17 | 6.15 | 4.73 | -1176 | Undruggable |
| 18 | 6.05 | 4.69 | -1319 | Undruggable |
| 19 | 5.84 | 4.62 | -1061 | Undruggable |
| 20 | 5.66 | 4.56 | -1187 | Undruggable |
| 21 | 5.44 | 4.49 | -1369 | Undruggable |

**Table S4:**
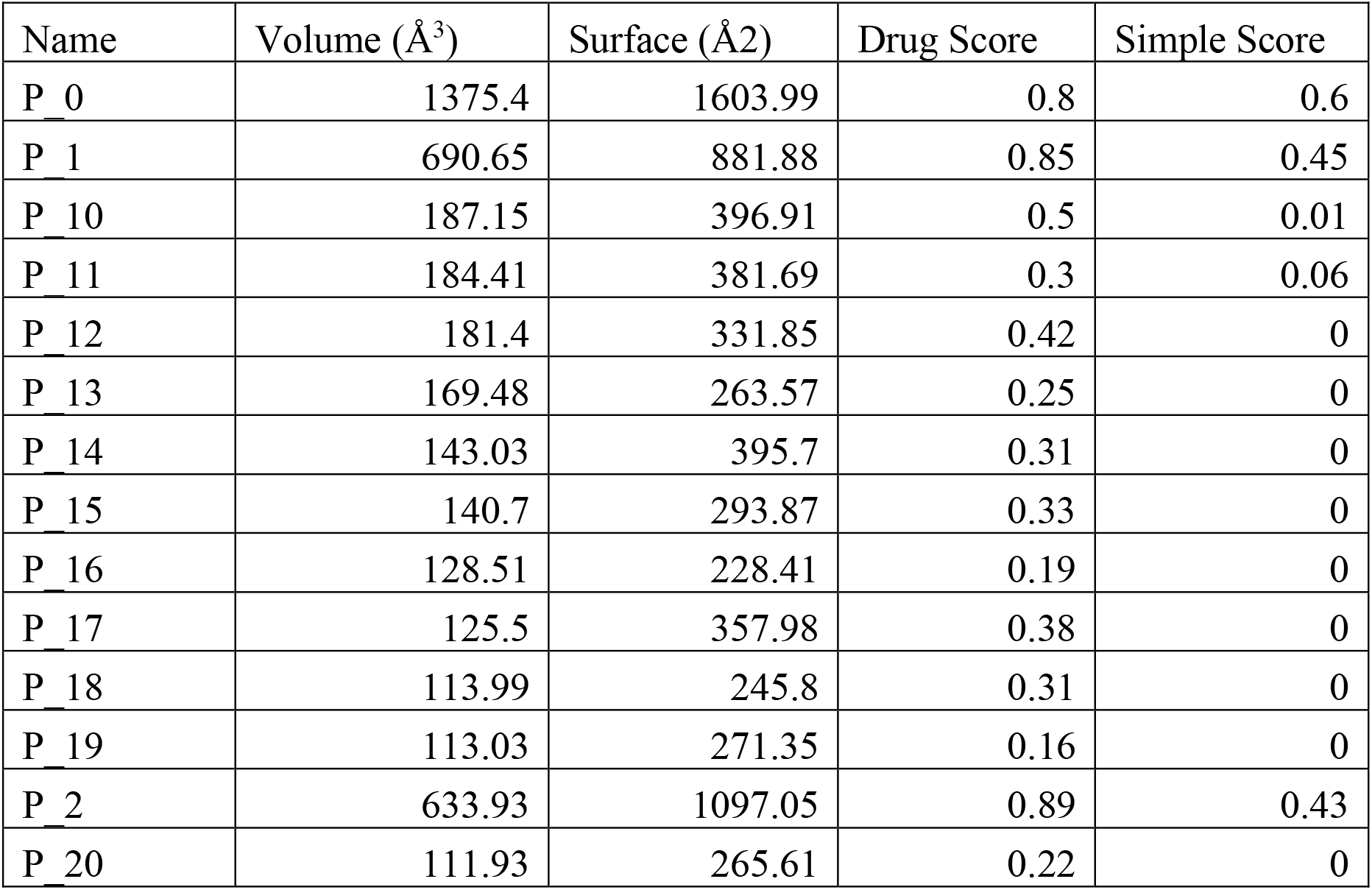
Results from the drug binding pocket prediction webserver DoGSitescorer.

**Figure S1.**
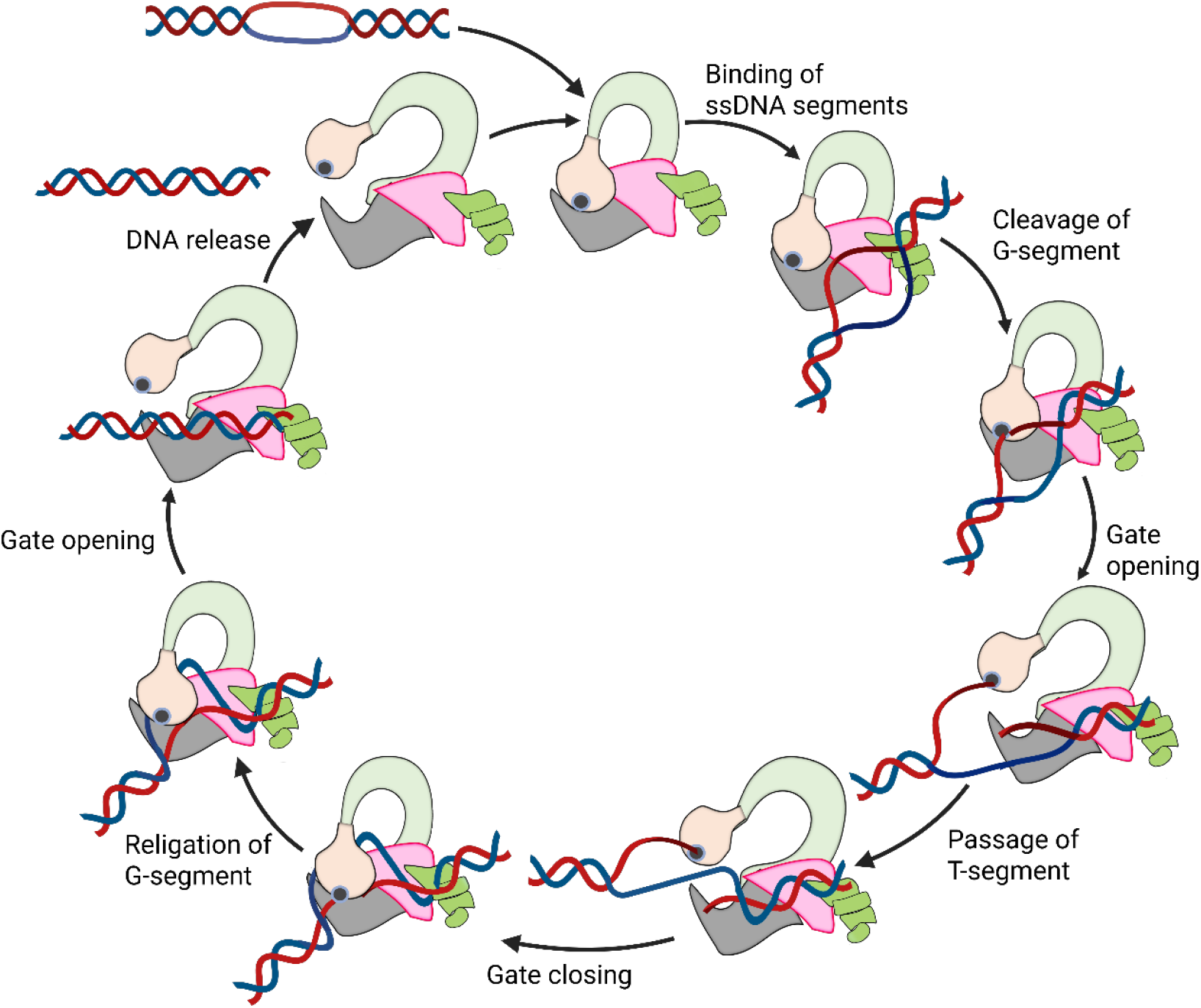
Model of catalytic cycle resulting in relaxation of negatively supercoils by *Mycobacterium tuberculosis* topoisomerase I.

**Figure S2.**
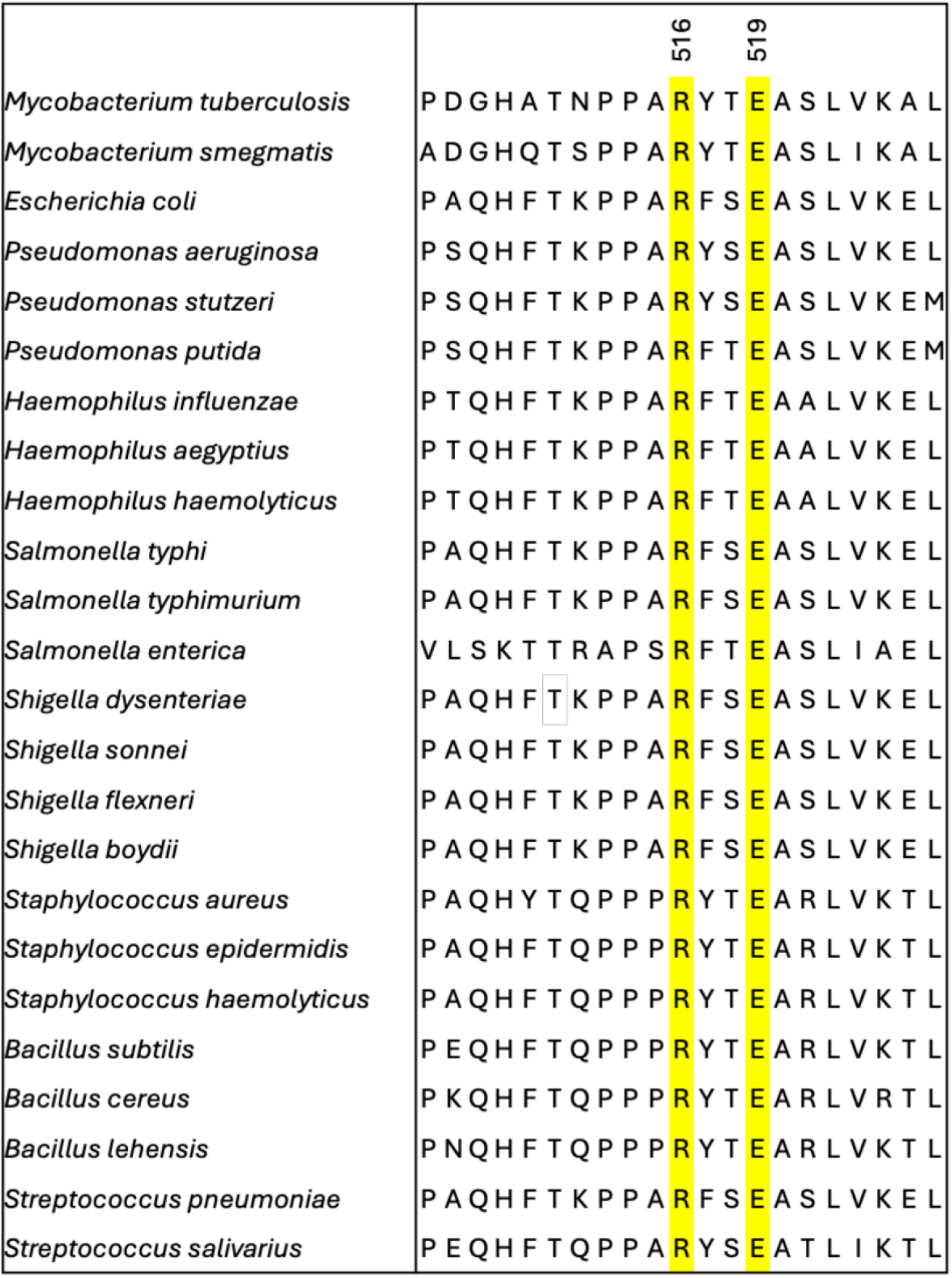
Alignment of bacterial topoisomerase IA sequences showing conservation of residues in predicted hinge region.

**Figure S3:**
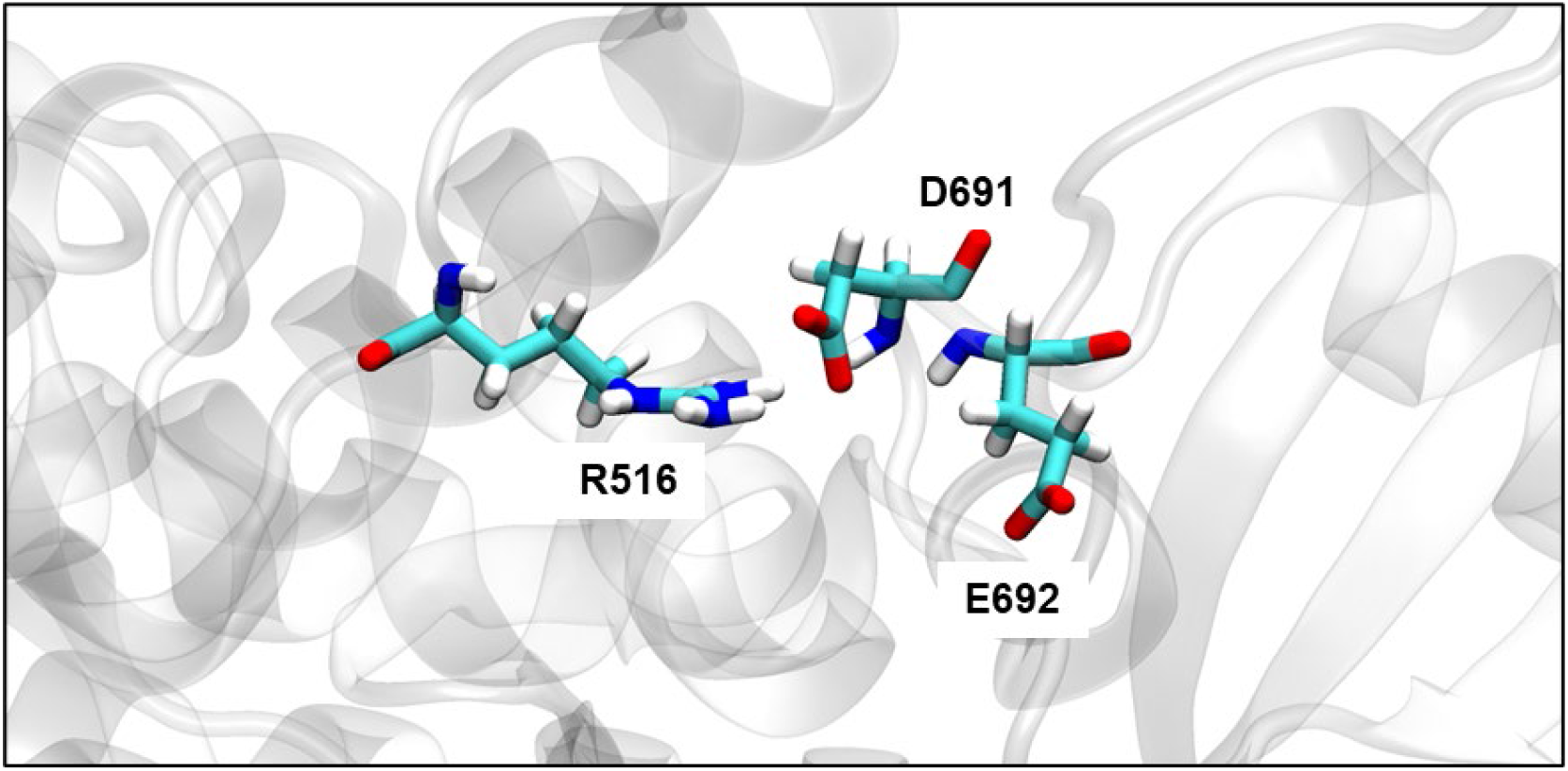
Figure showing the proximity of a negatively charged E692 to R516, to potentially compensate for the negative charge loss in the D691A mutation

